# Current limitations in predicting mRNA translation with deep learning models

**DOI:** 10.1101/2024.01.18.576214

**Authors:** Niels Schlusser, Asier González, Muskan Pandey, Mihaela Zavolan

**Affiliations:** Biozentrum, University of Basel, Spitalstrasse 41, Basel, 4056, Switzerland

**Keywords:** Translation control, Deep learning, Explainable AI, Systems biology

## Abstract

**Background:** The design of nucleotide sequences with defined properties is long-standing problem in bioengineering. An important application is protein expression, be it in the context of research or the production of mRNA vaccines. The rate of protein synthesis depends on the 5’ untranslated region (5’UTR) of the mRNAs, and recently, deep learning models were proposed to predict the translation output of mRNAs from the 5’UTR sequence. At the same time, large data sets of endogenous and reporter mRNA translation have become available.

**Results:** In this study we use complementary data obtained in two different cell types to assess the accuracy and generality of currently available models of translation. We find that while performing well on the data sets on which they were trained, deep learning models do not generalize well to other data sets, in particular of endogenous mRNAs, which differ in many properties from reporter constructs.

**Conclusions:** These differences limit the ability of deep learning models to uncover mechanisms of translation control and to predict the impact of genetic variation. We suggest directions that combine high-throughput measurements and machine learning to unravel mechanisms of translation control and improve construct design.

## 1 Background

The translation of most mRNAs into proteins is initiated by the recruitment of the eIF4F complex at the 7-methylguanosine cap, followed by eIF3, the initiator tRNA and the 40S subunit of the ribosome [1]. The 40S subunit scans the mRNA’s 5’ untranslated region (5’UTR) until it recognizes a start codon, then the 60S subunit joins to complete the ribosome assembly and initiate protein synthesis. Initiation is the limiting step of translation, largely determining the rate of protein synthesis [2]. It is influenced by multiple features of the 5’ untranslated region (5’UTR), from the structural accessibility of the cap-proximal region [3], to the strength of the Kozak sequence around the start codon (consensus gccRccAUGG, upper case - highly conserved bases, R = A or G, [4]), and the number and properties of upstream open reading frames (uORFs), that can hinder ribosome scanning to the main ORF (mORF), inhibiting its translation [5–8]. These (and presumably other) factors lead to initiation rates that differ up to 100 fold between mRNAs [9], and a similarly wider range of protein relative to mRNA abundance [10].

Accurate prediction of protein output from the mRNA sequence is of great interest for protein engineering and increasingly relevant with the rise of RNA-based therapies. This has prompted the development of both experimental methods for the high-throughput measurement of protein outputs as well as of computational models that can be trained on these data. An important development has been the introduction of ribosome footprinting, a technique for capturing and sequencing the footprints of translating ribosomes (RPFs) on individual mRNAs [2]. The ratio of normalized RPFs and RNA-seq reads over the coding region is used as an estimate of ”translation efficiency” (TE), which is considered a proxy for the synthesis rate of the encoded protein [2]. Ribosome footprinting has been applied to a variety of cells and organisms [11], yielding new mechanistic and regulatory insights (e.g. [12, 13]). An early study of yeast translation concluded that up to 58% of the variance in TE can be explained with 6 parameters, though the most predictive was the mRNA level expression of the gene, which is not a feature that can be derived from the sequence of the mRNA [6]. At the same time, massively parallel reporter assays (MPRA) were developed to measure translation for large libraries of reporter constructs, further used to train deep learning (DL) models. A convolutional neural network (CNN) [14] explained 93% of the variance in the mean ribosome load (MRL) of reporter constructs, but less, 81%, for 5’UTR fragments taken from endogenous mRNAs. The CNN also recovered some of the important regulatory elements such as uORFs [14]. More recently, a novel experimental design was used to accurately measure the output of yeast reporters driven by natural 5’UTRs [15], while novel DL architectures and training approaches aimed to improve prediction accuracy [16, 17]. Potential limitations of DL models built from synthetic sequences is that it is *a priori* unclear whether the training set contains the regulatory elements that are relevant *in vivo* and whether the features extracted by the model generalize well across systems such as cell types and readouts of the process of interest. These bottlenecks limit not only the understanding of regulatory mechanisms but also the use of derived models for predictions of functional impact and for construct design. To assess whether these issues impact the current RNA sequence-based models of translation, we carried out a detailed comparison of model performance in a standardized setting that uses complementary data sets obtained in two distinct cell types. We trained and applied models to the prediction of translation output in yeast and human cells, addressing the following questions: (1) are models trained on synthetic sequences able to predict the translation output of endogenous mRNAs in the same cellular system? (2) do these models generalize between different cellular systems (different cell types, different species)? (3) what is their parameter-efficiency (fraction of explained variance per model parameter)? (4) what are the conserved regulatory elements of translation that have so far been learned by DL models?

### 1.1 Experimental measurements of translation output

The current method of choice for measuring the translation output of endogenous mRNAs is ribosome footprinting, consisting in the purification and sequencing of mRNA fragments that are protected from RNase digestion by translating ribosomes [2]. The TE of an mRNA is then estimated as the ratio of ribosome-protected fragments (RPFs) obtained from the mRNA by ribosome footprinting and coding-region-mapped reads obtained by RNA-seq from the same sample [18]. Ribosome footprinting has been applied to many systems, including yeast cells [6] and the human embryonic kidney cell line HEK 293 [19], for which a variety of omics measurements are available. Importantly, MPRA of translation were carried out in these cell types, giving us the opportunity to determine whether reporter-based models can predict translation of endogenous mRNAs in a given cell type. Figure 1 summarizes the main approaches used to measure translation in yeast and human cells, starting from the just-described ribosome footprinting technique (Fig. 1A). The MPRA approach used by [14] to generate the Optimus50/100 MPRA data sets (Fig. 1B) consists in the transfection of *in vitro*-transcribed mRNAs with randomized 5’UTRs upstream of the eGFP coding region into HEK 293 cells, followed by sequencing of RNAs from polysome fractions. The MRL, i.e. the average number of ribosomes on individual mRNAs is derived from abundance profile of individual mRNAs along polysome fractions. In another approach, called DART (for direct analysis of ribosome targeting), Niederer and colleagues [15] have synthesized *in vitro* translation-competent mRNAs consisting of natural 5’UTRs and a very short (24 nucleotides (nts)) coding sequence. A few mutations were introduced in the 5’UTRs, as necessary to unambiguously define the translation start. After incubation with yeast extract, ribosome-associated mRNAs were isolated, sequenced, and a ribosome recruitment score (RRS) of an mRNA was calculated as the ratio of its abundance in the ribosome-bound fraction relative to the input mRNA pool (Fig. 1C). A previously-developed MPRA used plasmids containing randomized 5’UTRs placed upstream of the *HIS3* gene to transform yeast cells lacking a native copy of the gene [20]. The amount of HIS3 protein generated from the reporters was assessed in a competitive growth assay, by culturing the yeast cells in media lacking histidine, and using the enrichment of reporter constructs in the output vs. the input culture as a measure of His3 expression (Fig. 1D).

**Fig. 1.**
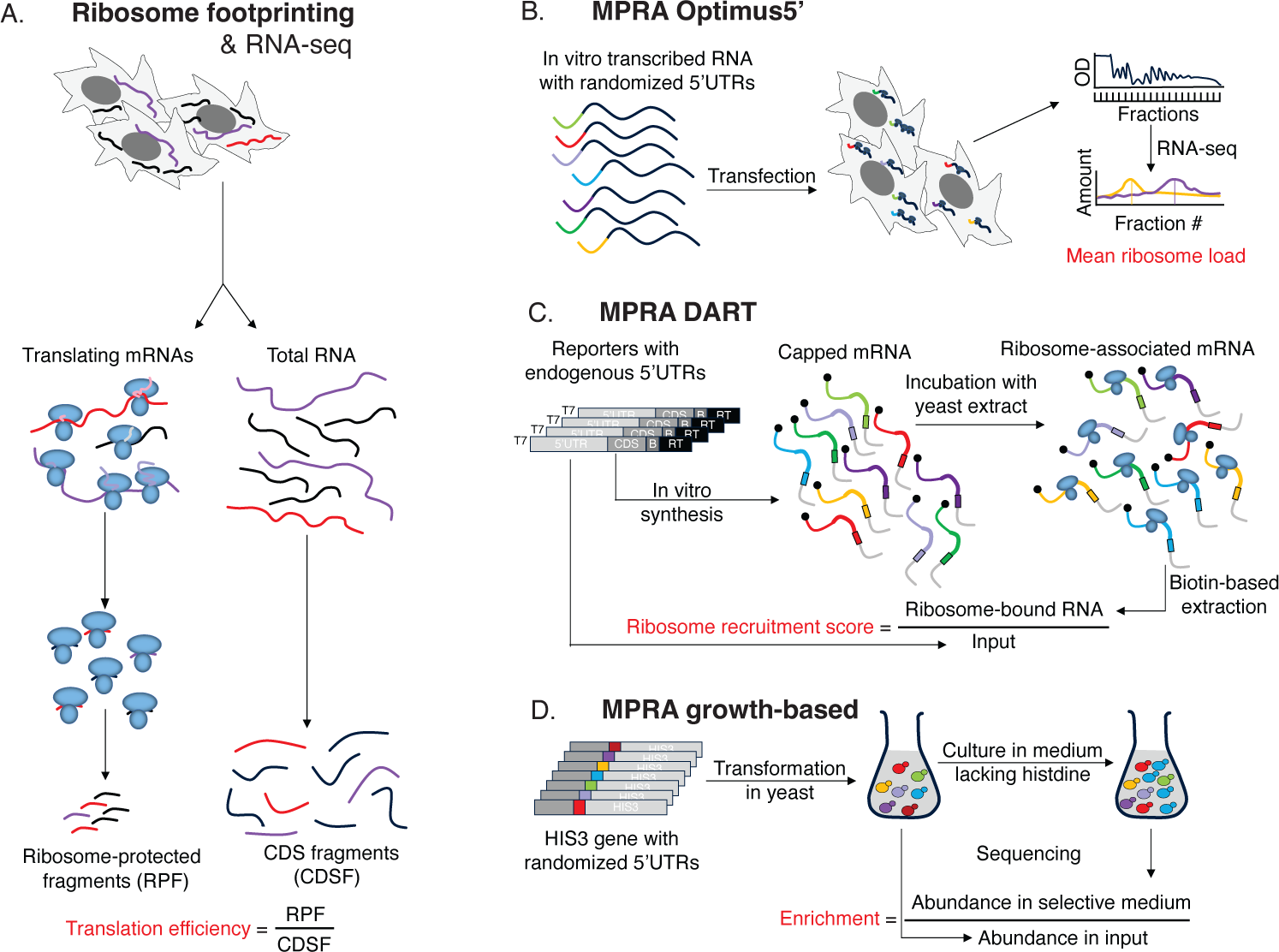
Experimental approaches to quantifying translation output. Sequencing of total mRNA and ribosome-protected fragments of endogenous mRNAs is used to estimate the translation efficiency per mRNA (A). Massively parallel reporter assays (MPRA) measure the output of constructs consisting in randomized 5’UTRs sequences attached to the coding region of a reporter protein (B). Sequencing of polysome fractions enables the calculation of a mean ribosome load per construct, which is used as a measure of translation output. DART (C) follows a similar approach with endogenous 5’UTRs, once upstream AUGs (uAUGs) located in the 5’UTR are mutated to AGU to avoid ambiguity in translation start sites. In an alternative MPRA in yeast, the enrichment of 5’UTRs driving expression of a protein required for growth served as proxy for the translation output of the respective constructs (D). More details can be found in the Extended Methods, Subsec. 1.1.

The reproducibility of experimental measurements sets an upper bound on the accuracy of models’ predictions of different types of data. For MPRA data sets the *R*^2^ of replicate measurements is usually very high, values of 0.95 being generally reported [14]. In contrast, the reproducibility of TE, which is a ratio of two variables - ribosome footprints and RNA-seq fragments mapping to a given mRNA -, is generally lower. In the HEK 293 ribo-seq data set that we analyzed [19], the *R*^2^ for RPFs was in the range 0.77 *−* 0.82, while for mRNA-seq 0.96, leading to *R*^2^ of TE estimates 0.47 *−* 0.52 (Supplementary Fig. 1). We further obtained an additional ribo-seq data set from another human cell line, HepG2, with the aim of exploring the limits of replicate reproducibility of this type of measurement and evaluating the conservation of TE between cell types, which is also important when applying a model trained in a particular cell type to predict data from another cell type. The TE estimates from HepG2 cells were more reproducible, with *R*^2^ for replicates 0.68*−*0.8. When comparing the TE estimates from HEK 293 and HepG2 cells we obtained an *R*^2^ = 0.31, which would be an upper bound on the accuracy of a model trained on one of these data sets in predicting the TEs in the other cell line.

To ensure comparability of our results with those of previous studies, we aimed to replicate their model training strategy, which generally involved setting aside the highest quality data (constructs with the largest number of mapped reads) for testing and using the rest for training [14, 16]. High expression is not the only determinant of measurement accuracy for endogenous mRNA data sets. For example, in yeast, the integrity of the sequenced RNAs was previously identified as key source of noise for TE estimates [6]. A proxy for RNA integrity is the transcript integrity score (TIN) [21], which quantifies the uniformity of coverage of the RNA by sequenced reads and ranges from 0 (3’ bias) to 100 (perfectly uniform coverage). As the TE reproducibility in HEK 293 cells increased with the TIN score (*R*^2^ = 0.67 *−* 0.75 for TIN *>* 70 vs. 0.47 *−* 0.52 for all), we used the mRNAs with TIN *>* 70 (*≈* 10% mRNAs) for testing and all the others for training.

### 1.2 Models for predicting translation output from the mRNA sequence

To explain the translation efficiency estimated by ribosome footprinting in yeast, [6] proposed a simple, 6-parameter linear model with the following features: lengths of the mORF and 5’UTR, G/C content of the 5’UTR, number of uAUGs, free energy of folding of the 5’cap-proximal region and the mRNA abundance. This linear model was surprisingly accurate (*R*^2^ = 0.58) in predicting the efficiency, though leaving out the mRNA level reduced the *R*^2^ to 0.39. Here we use a similar model as baseline to assess the parameter efficiency of DL models, i.e. the fraction of explained variance per model parameter. The features of our linear model are: we use the same length and G/C content measures, the 5’UTR folding free energy divided by the 5’UTR length, the number of out-of-frame upstream AUGs (OOF uAUGs), the number of in-frame upstream AUGs (IF uAUGs), and the number of exons in the mRNA [22]. A bias term adds an additional parameter.

The first type of DL architecture trained on MPRA data was the Optimus5’ CNN [14], operating on one-hot-encoded 5’UTR sequences. Optimus5’ has 3 convolutional layers, each with 120 filters of 8 nucleotides (nts), followed by two dense layers separated by a dropout layer. The output of the last layer is the predicted translation output, i.e. the MRL for the HEK 293 cell line data set (Fig. 1B, 2A) and the relative growth rate for yeast cells [20] (Fig. 1D). While reaching very good performance in predicting the MRL for synthetic sequences, Optimus5’ could only make predictions for UTRs of up to 100 nts, which account for only 32% of annotated human 5*^′^*UTRs. Longer 5’UTRs could be accommodated by truncation to the upstream vicinity of the start codon. The Optimus5’ model just described has 474’681 parameters.

The 5’UTR length limitation was overcome by Framepool [16], another CNN containing 3 convolutional layers with 128 7-nts filters (Fig. 2B). Importantly, Framepool slices the output of the third convolutional layer according to the frame relative to the start codon, then pools the data frame-wise, taking the maximum and average values for each pool. This allows both the processing of sequences of arbitrary length and the detection of motifs in specific reading frames. The frame-wise results are the input for the final two dense layers. For variable length 5’UTRs, Framepool was shown to yield somewhat improved predictions relative to Optimus5’ [16], with a smaller number of parameters, 282’625.

**Fig. 2.**
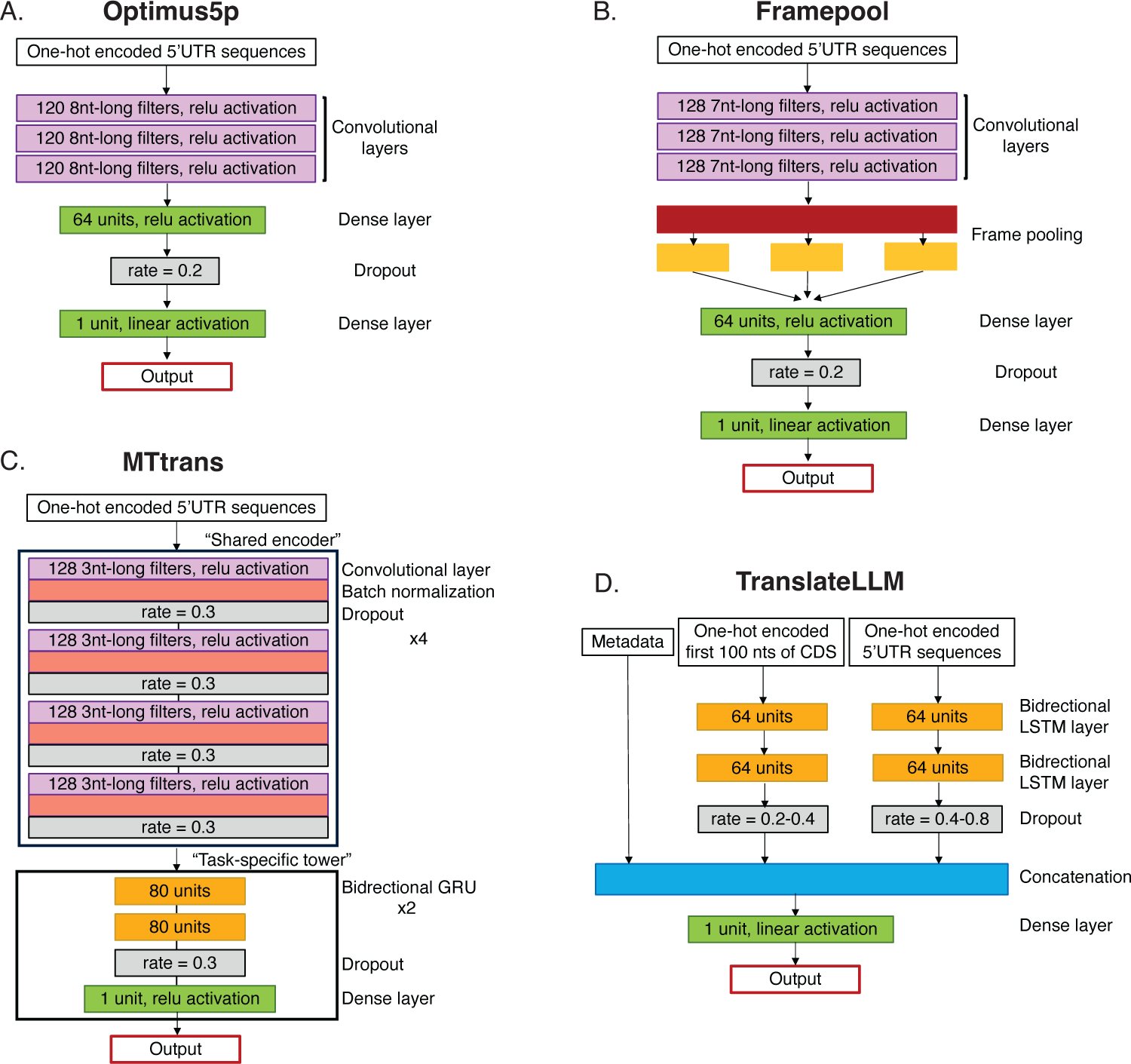
Architectures of different artificial neural networks used to predict the output of translation. Optimus5’ [14] uses three convolutional layers and two dense layers (A), Framepool (B) similar, but with a customized frame-wise pooling operation between convolutional and dense layers [16], MTtrans (C) stacks a ”task-specific” tower of two recurrent and one dense layers on top of a ”shared encoder” of four convolutional layers. TranslateLLM (D) consists of three sub-networks: a two-layer bidirectional LSTM network for the 5’UTRs, another two-layer bidirectional LSTM network for the first 100 nts of the ORF, and non-sequential input features previously found to control translation. For further information, we refer to the Extended Methods section, Subsec. 1.3.

MTtrans [17] is the most recently proposed DL model (Fig. 2C). Its basic premise is that the elements controlling translation generalize across data sets generated with different experimental techniques. Each data set is viewed as a task. The model combines 4 convolutional layers with batch normalization, regularization and dropout, with two bidirectional gated recurrent unit (GRU) layers and a final dense layer. GRUs are recurrent layers that process the input sequence token by token and map these to an internal space that allows contextual information to be preserved. Inputs of different lengths can naturally be processed by this architecture. The 4 convolutional layers, which the authors called ’shared encoder’, are assumed to be universal among different prediction tasks, while the recurrent and dense layers (’task-specific tower’) are specific to each task. The shared encoder is therefore trained on multiple data sets, while the task-specific tower is trained only on the respective task. In comparison to Optimus5’ and Framepool, MTtrans provides an increase in *R*^2^ of 0.015 *−* 0.06 in prediction accuracy, depending on the data set [17]). Interestingly, training MTtrans on multiple data sets at once rather than in a sequential, task-specific manner, achieved an almost similar effect. While we were able to obtain the code for MTtrans from [23], we were unable to run the code ”out-of-the-box”. Therefore, we set up MTtrans as described in the conference publication [17] though this left many details unclear, i.e. the exact layout of the ”task-specific tower”, its recurrent and dense layers, the number of training epochs, the exact training rate schedule, and criteria for early stopping. It also led to a different number of parameters in our implementation, 776’097, compared to the number reported by the authors 2.1 million. Consequently, we trained with a callback that automatically stops once overfitting is reached and restores the best weights. Although, in our experience, these are details have only a minor impact on the model performance, we note that our results differ to some extent from those reported in [17]. The use of GRUs in the task-specific tower allows MTtrans to predict output for any 5’UTR length.

While DL models become increasingly more parameter-rich, their performance improves only marginally, leading to a decrease in the gained accuracy per parameter. We were therefore interested in whether the parameter-efficiency of DL models can be improved, i.e. whether the top performance can be achieved with smaller rather than larger models. To address this, we turned to long short-term memory networks (LSTMs), a variety of recurrent neural networks (RNNs) designed to detect and take advantage of long-range dependencies in sequences [24]. While such dependencies are expected in structured 5’UTRs, LSTMs have not been applied yet to the prediction of translation output. We therefore implemented here two LSTM-based architectures: one operating only on 5’UTR sequences and a second one, TranslateLLM, operating not only on 5’UTRs, but also on the first 100 nts of the associated coding regions and the non-sequential features of the linear model described above. The extended TranslateLLM allows for factors such as the secondary structure and codon bias in the vicinity of the start codon [5] to impact the translation output. One-hot-encoded sequences are fed into two bidirectional LSTM layers, the outputs of the second layers are concatenated and sent to dense layer which predicts the output (Fig. 2D). TranslateLLM has 268’549 - 268’552 parameters, while the 5’UTR-only LSTM model has 134’273 parameters. We further note that, depending on the experimental design, not all data sets to which a given model is applied require the same number of parameters. For instance, a data set in which all sequences have the same length like Optimus50 does not require the sequence length as a parameter in TranslateLLM or the linear model. Similarly, as the first 100 nts of the ORF are the same in all MPRA data sets, the associated parameters are not needed in TranslateLLM, which reduces the number of parameters to about 50% relative to the full model.

## 2 Results

### 2.1 Available DL models do not generalize well across experimental systems

The results of our comprehensive tests of prediction accuracy of all models across multiple data sets are summarized in Fig. 3A. The most salient result is that differences in performance between DL models applied to a particular data set are much smaller than differences between applications of the same model to distinct data sets. In particular, DL models can be trained on synthetic constructs to predict the output of leave-out constructs, but they cannot be trained well on TE data to predict the translation of endogenous mRNAs (compare lines 1,2,3 and 4,5 in Fig. 3A). Fig. 3B shows scatter plots of ribosome load predicted by each of the discussed DL architectures against their measured counterparts. It can be clearly seen that OOF uAUGs are strongly inhibiting translation. Moreover, the TranslateLLM predictions are most uniformly spread around the diagonal. The size of the training data set is not strictly a limiting factor, because DL models can be trained to some extent on the relatively small DART data set of *∼* 7*^′^*000 natural yeast 5’UTRs (Fig. 3A, l. 6). Furthermore, models trained on synthetic 5’UTRs do not predict the TE of endogenous mRNAs measured in the same cell type (see Fig. 3A ls. 7,8). This reduced performance was previously attributed to the different experimental readout and to the influence of the coding sequence, which is the same in MPRA, but different in TE assays [16]. To test this, we applied the models trained on human MPRA data to the prediction of MPRA data from yeast and vice versa. This involved not only a very different readout of translation (MRL in human, growth-dependent enrichment of 5’UTRs in yeast), but also entirely different organisms. In both cases, the cross-system performance substantially higher, *R*^2^ = 0.41 and 0.64 (c.f. Fig. 3A ls. 9,10), compared to the performance of the model trained on synthetic data in predicting the TE data in the same cell type. Thus, the type of readout is not the main factor behind the reduced predictability of TE data. Another limiting factor, not discussed before, could be the accuracy of the experimental measurements. MPRA and DART-based measurements are very reproducible, with *R*^2^ 0.95, while the TE estimates much less so (*R*^2^ *≈* 0.5 for the HEK 293 data set, Supp. Fig. 1). Thus, the TE data may be less predictable as it is also more noisy. However, the measurement accuracy is not a factor in the highly reproducible DART experiments, yet models trained on synthetic construct data from yeast could not predict the RRS measured in the DART experiment, also done in yeast. Altogether, these results indicate that synthetic sequences differ substantially from natural, evolved, 5’UTRs, leading to models trained on synthetic data not being able to adequately capture 5’UTR features that are relevant for the translation of endogenous mRNAs. We also applied a transfer-learning strategy to human HEK 293 data, where we first trained the models on the Optimus100 data set, then re-trained the last layer on endogenous data, and finally some epochs of training the entire network on the endogenous data. The results are displayed in Fig. 3A l. 13. Applying transfer learning indeed lead to a small performance increase of 0.04 in *R*^2^.

**Fig. 3.**
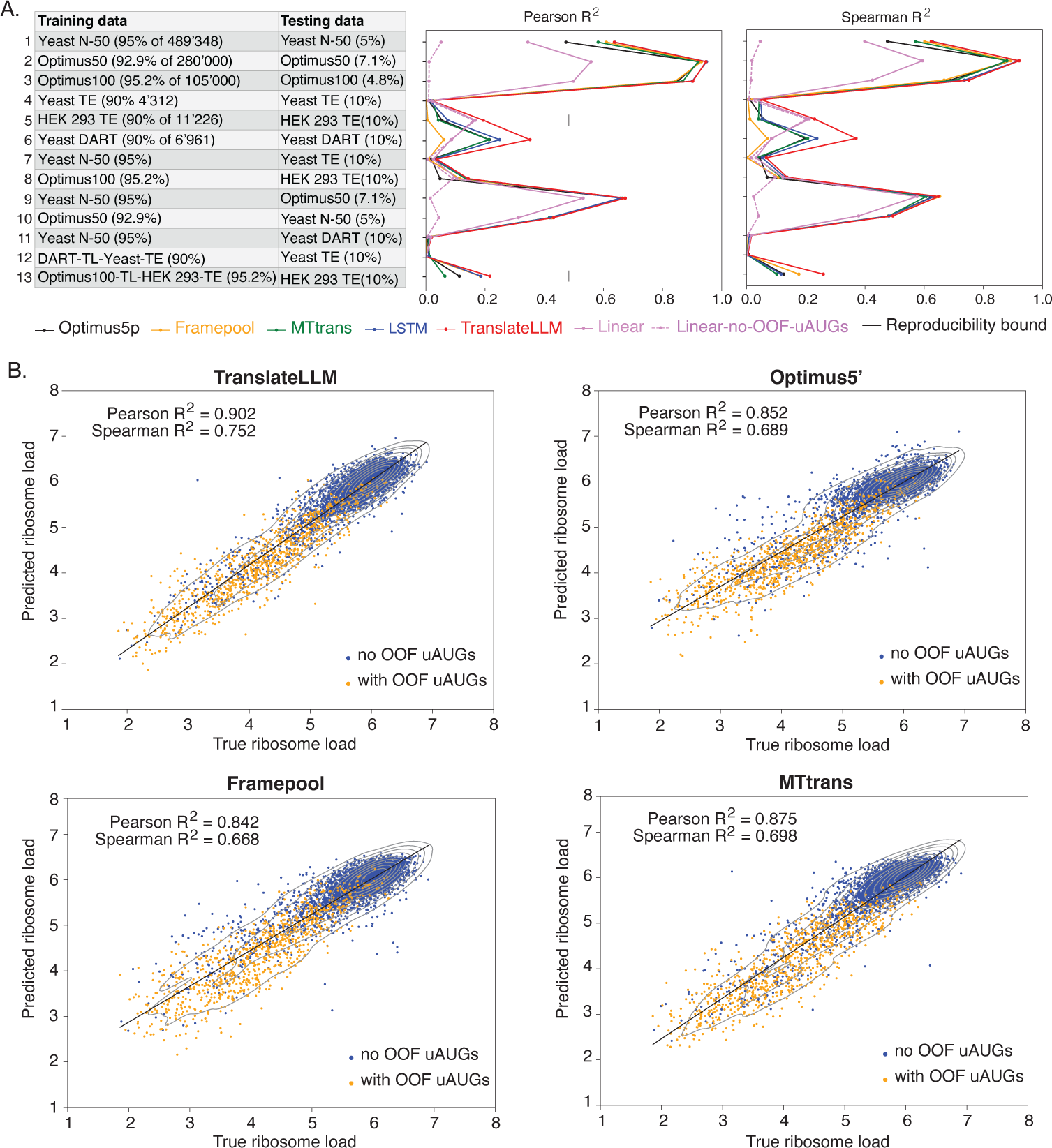
Performance of all evaluated models in different application scenarios (A), measured by the Pearson and Spearman correlation coefficients 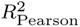 and 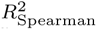 between experimentally measured and predicted translation output in the test data. Different random splits of the DART data lead to variations in *R*^2^ of ≲ 0.02, with the exception of the Framepool model, which had differences of up to 0.16 between splits. (B) Correlation of predicted and true ribosome load of four model architectures trained on the Optimus100 data set. OOF uAUGs clearly inhibit translationinitiation. TranslateLLM predicts the most even scattering pattern around the diagonal.

As seen above, the yeast-human cross-species prediction accuracy is substantial, indicating that the translation regulatory elements inferred from synthetic constructs in the two species are partially conserved. Given that the cell type in which a model is developed will generally different from the cell type where model predictions are of interest, we asked whether the TE of human mRNAs are largely similar across human cell lines. We thus generated ribosome footprinting data from the human HepG2 liver cancer cell line and compared the TE inferred from this cell line with those from the HEK 293 cell line data set. The estimates of TE were more reproducible than those from the HEK 293 cells (*R*^2^ = 0.68 *−* 0.8 for replicate experiments). The TEs estimated from HEK 293 were moderately similar to those from HepG2 cells (*R*^2^ = 0.31), especially when considering mRNAs with TIN *>* 70 (*R*^2^ = 0.44). This indicates that within an organism, transcript-intrinsic properties contribute substantially to the variation in translation output relative to the cellular context. This is a good basis for developing models in model systems, provided that the protocol allows for highly accurate measurements on translation output (Supplementary Fig. 2).

Although DL models are not generally benchmark against simple models with more limited predictive power, this test provides an assessment of parameter-efficiency (gain in predictive power per parameter), as well as insights into model interpretation. Trained on synthetic construct data, the 8-parameter linear model described above could explain as much as 60% of the variance in the respective test sets, which is quite remarkable given the size of the model. In addition, this model could also be trained to some extent on TE measurements of endogenous mRNAs. Strikingly, the accuracy of cross-system predictions of synthetic construct-based DL models is similar to the accuracy of linear models inferred from the respective data sets. This indicates that the conserved mechanisms of translation control learned by the DL architectures are represented in a small number of features, and that currently available DL architectures are heavily over-parameterized.

### 2.2 Reporter sequences differ in translation-relevant properties from endogenous mRNAs

To further identify the most predictive and conserved features, we inspected the weights learned by the linear model from individual data sets (Supp. Fig. 4A). We found that only the uAUGs, especially those located out-of-frame (OOF) with respect to the mORF, consistently contributed to the prediction of translation output across all systems. OOF uORFs/uAUGs are known to repress translation, by hindering ribosome scanning towards the mORF [25] and triggering the mechanism of non-sense mediated mRNA decay [26]. uAUGs contribute much more to the translation output of human or yeast reporters constructs compared to endogenous mRNAs, which is a reflection of differences in sequence composition between synthetic and natural 5’UTRs (Supplementary Fig. 3). This suggests that models trained on synthetic sequences will incorrectly weigh the translation-relevant features they learned from these sequences when predicting the output of natural 5’UTRs, leading to reduced prediction accuracy. To illustrate this, we carried out a simulation using the Optimus50 data set: we set aside the 20’000 constructs with highest coverage in mRNA sequencing for testing as before, but trained the Optimus5’ model on the subset of remaining constructs that did contain uAUGs. As shown in Supp. Fig. 4B, the resulting model performs poorly on the test set, specifically on the subset of test sequences that do contain uAUGs. In addition, even this model, that could, in principle, learn other regulatory elements of translation, does not predict the translation output of the DART dataset of natural yeast 5’UTRs lacking uAUGs, see l. 11 of Fig. 3A. These results demonstrate that the similarity of distributions of translation-relevant features among training and test set are key to the ability of the DL model to generalize. Having undergone extensive selection under a variety of constraints, endogenous 5’UTRs likely accumulated multiple elements that control their translation, elements that are probably not represented among synthetic 5’UTRs. This leads to large differences in performance when models trained on synthetic data are applied to other data sets.

Previous studies reached different conclusions concerning the impact of IF uAUGs on translation [15, 20, 27, 28]. To clarify this, we determined the relationship between the location of OOF and IF uAUGs in the 5’UTR and the translation output of the mRNAs, in both yeast and human sequences, synthetic or endogenous. To avoid a superposition of effects from multiple uAUGs, we analyzed only constructs with a single uAUG in the 5’UTR. As shown in Supp. Fig. 4C-H, the repressive effect of OOF uAUGs on the translation of synthetic constructs is largely position-independent, while the repressive effect of IF uAUGs increases with their distance from the mORF. Similar trends are apparent when we use all constructs, computing the distance-dependency for only the 5’-most uAUG (Supp. Fig. 4E-H). Endogenous mRNAs show the same trends, though less pronounced Supp. Fig. 4G-H. These results indicate that both the frame and the distance of uAUGs with respect to the mORF should be taken into account when predicting their impact on translation.

### 2.3 A more accurate and parameter-efficient DL model to predict the impact of 5’UTR sequence variation on translation

To provide a more accurate model of endogenous mRNA translation, accommodating different constraints on uAUGs and improving parameter-efficiency, we turned to LSTM-based architectures. The two architectures that we implemented, LSTM and TranslateLLM (see Fig. 2) performed similarly on the synthetic data sets, and were more accurate that the other DL models tested. The largest performance gain was reached for RNAs with IF uAUGs, as may be expected from the model’s treatment of sequence context (Supp. Fig. 5). The similar performance of LSTM and TranslateLLM on synthetic data indicates the LSTM can learn correlates of the non-sequential features represented in TranslateLLM. However, these features were important for the performance of TranslateLLM on the endogenous HEK 293 TE data (Fig. 3A and Supp. Fig. 4A).

To demonstrate the relevance of DL models for interpreting the functional significance of single nucleotide polymorphisms (SNPs), Sample et al. [14] measured the MRL of constructs with 50 nts-long fragments of natural 5’UTRs as well as of variants with naturally-occurring SNPs. TranslateLLM predicted better the measured MRL of these sequences than Optimus5’ model (Fig. 4A-B). However, in this experiment, 5’UTR sequences were taken out of their endogenous context, which, as we have shown above, is important for the prediction of translation output and thereby functional impact. Therefore, we sought to improve the prediction of SNP effects on translation taking advantage of the insights provided by our analyses. We used transfer learning (TL) to extract information from both synthetic and endogenous 5’UTRs and we applied the resulting model to all the 5’UTR-located SNPs from the ClinVar database [29, 30], in their native 5’UTR context. 84’128 of the 2’300’005 SNPs were located in 5’UTRs and of these, 7’238 were located in mRNA isoforms (one per gene) expressed and with measured TE in HEK293. As shown in Fig. 3A l. 13 the TL strategy leads to better predictions than the training on endogenous data alone, and also better than the predictions of other DL models trained by TL. The distribution of log-ratio of predicted translation output of variant and wildtype sequences is shown in Fig. 4C. 110 of the 7238 variants are predicted to affect the TE by 10-fold or more, 34 increasing and 76 decreasing the TE compared to the wildtype sequence. Interestingly, despite the large predicted impact, none of the 110 SNPs create or destroy an uAUG. However, overall, while absolute numbers of uAUG changes are small (328 of 7’238 variants), creation/destruction of an uAUG was associated with a predicted reduction/increase of translation output. Moreover, the pathogenic variants had a small bias for increased TE (Fig. 4D).

**Fig. 4.**
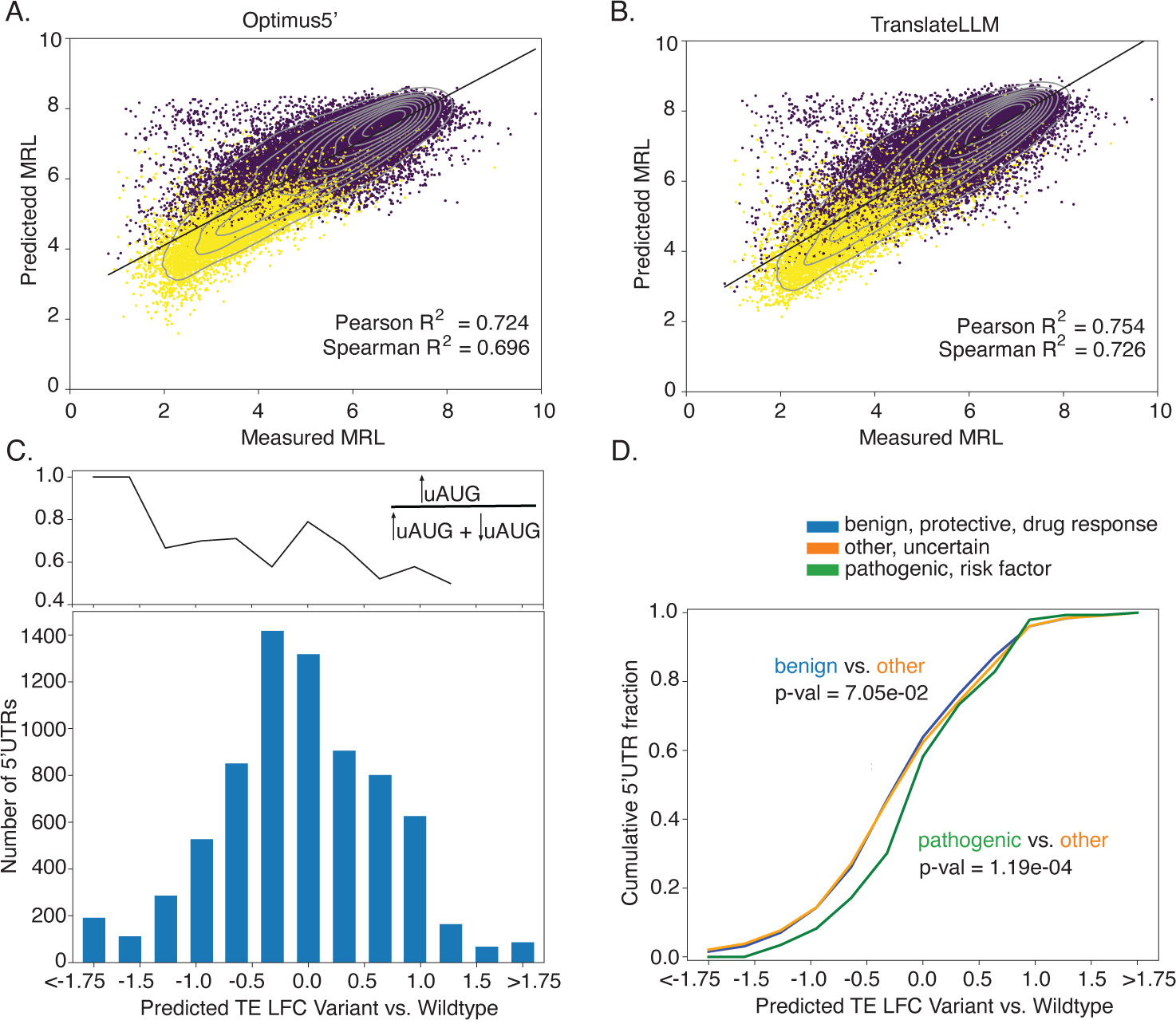
Effect of 5’UTR-located variants on mRNA translation output. (A) Optimus5’ was trained on a pool of randomized 50nt long sequences and applied to a pool of equally long known variants (see Subsubsec. 5.2.3). Yellow points indicate 5’UTRs with OOF uAUGs, purple points without OOF uAUGs. Same was done for the TranslateLLM architecture in panel (B). We use TranslateLLM to predict the TE of known clinical variants of endogenous sequences from the ClinVar database [29]. We compare the predictions to their (measured) wildtype counterparts (see Subsubsec. 5.2.11) and obtain log-fold changes of the translation efficiency (TE LFC). It follows a normal distribution (C), where a negative TE LFC can be associated with a propensity for the variant to create uAUGs, while positive TE LFC is associated with a propensity of breaking uAUGs. Clinical variants with known pathogenic phenotype (clinical significance annotation: pathogenic, likely pathogenic, risk factor) are predicted to significantly increase the TE compared to their neutral counterparts (D), whereas variants with benign phenotypes (clinical significance annotation: benign, likely benign, protective, drug response) do not significantly alter the distribution, as demonstrated by Kolmogorov-Smirnov tests.

## 3 Discussion

The wider dynamic range of protein compared to mRNA expression suggested an important role of translation control in determining protein levels [10]. Initiation is the limiting step of translation [2, 6, 7], modulated by a variety of regulatory elements in the 5’UTRs [6, 31], from uORFs to internal ribosome entry sites [32–34]. With the rise in mRNA-based therapies, the interest in designing 5’UTRs to drive specific levels of protein expression has surged [14], prompting the development of DL models to predict the translation output of mRNAs from the 5’UTR sequence. To satisfy the large data needs of these models, a few groups have devised innovative approaches to measure the translation output of large numbers of constructs, containing either random 5’ UTRs or fragments of endogenous sequences [14, 15, 20]. DL models trained on these data achieve impressive prediction performance on leave-out data, and are used to identify sequence elements that modulate translation, the most predictive among these being uAUGs/uORFs. However, DL models trained on synthetic data do not predict well the translation output of endogenous mRNAs. In this study we carried out an extensive comparison of models of translation across multiple data sets and settings, to understand the limits of their applicability and generality.

We took advantage of two systems in which the translation output has been measured for both synthetic and endogenous 5’UTRs, namely yeast [6, 20], and HEK 293 cells [14, 19]. For yeast, an additional library of *∼* 12*^′^*000 endogenous 5’UTRs devoid of uAUGs was tested for their ability to recruit ribosomes [15]. We observed the best performance in the yeast-human cross-prediction of translation output of synthetic constructs, even though the readouts of the assays were very different for the two organisms. This prediction relies on a small number of conserved determinants of translation output, in particular uAUGs, as underscored by the similar performance achieved with an 8-parameter linear model trained on the same data sets. However, models trained on synthetic constructs do not predict the translation output of endogenous mRNAs. The influence of the coding region or to trans-acting factors do not play a key role, as demonstrated with the various yeast data sets, where these factors were controlled. Rather, endogenous sequences have been selected in evolution under multiple constraints, not limited to translation output, and have acquired a variety of regulatory elements that are not well-represented in the randomized 5’UTRs. This leads to models trained on synthetic data not having the possibility to learn such features. We could most clearly demonstrate this with a simulation, in which a model trained on sequences lacking uAUGs performed poorly on a data set in which these elements are represented. While in this case the outcome may seem obvious, as uAUGs are important modulators of translation output, there are likely many other elements that are not well represented among synthetic sequences yet affect the translation output in various ways, including for e.g. by influencing the mRNA stability. All of these factors ultimately contribute to the poor performance of models trained in synthetic 5’UTRs on predicting the translation output of endogenous mRNAs.

The same issues likely compound the prediction of SNP effects. As the genetic variation of human populations is being mapped, DL models are increasingly used to predict various molecular phenotypes, including the translation output of mRNAs [14, 16]. Genetic variation is manifested in the native gene expression context, implying that predictions of models trained on synthetic sequences will not be reliable. Given that with TranslateLLM we were able to explain more of the variance in TE compared to other DL models, we also sought to provide updated predictions of the potential impact of ClinVar variants on TE [29]. Surprisingly, variants classified as pathogenic are predicted to more often increase than decrease the TE of the respective mRNA, i.e. they tend to be gain-of-function variants. Interestingly, the increase is not generally explained by the removal of a repressive uAUG, as relatively few SNPs changed the number of uAUGs in the 5’UTRs. These 1000 SNPs predicted to most increase the TE came from genes involved in protein and organelle localization (Suppl. Table 1), predictions that could be tested in a future study.

That *∼* 60% of the variance in MPRA data can be explained with models constructed from such distant species such as yeast and human indicates that the models have learned deeply conserved mechanisms of controlling the translation output. That the simple, 8-parameter linear model, performs almost on par with the DL models in this setting indicates not only that these mechanisms are reflected in a small number of mRNA features, but also that the DL models are heavily over-parameterized. Indeed, the cross-species prediction power comes largely from the OOF uAUGs, as demonstrated by the poor performance of the linear model lacking this element. The 5’UTR G/C content and/or free energy of folding appear to be additional conserved regulatory elements, with more prominent role in explaining the translation output of evolved 5’UTRs.

To the extent to which synthetic data sets and DL models are used to uncover molecular mechanisms, it is important to ponder whether the synthetic sequences cover these mechanisms, as well as whether the model architecture allows for the appropriate representation of these mechanisms. This is, of course, difficult to ensure a priori, when the mechanisms are unknown. However, an improved grasp of the parameter efficiency of models and their interpretation should facilitate the discovery of regulatory mechanisms and avoid false inferences. For example, CNN type of architectures may be able to encode correlates of RNA secondary structure sufficiently well to predict the translation of short, synthetic 5’UTRs. Yet the sequence motifs learned by the CNN need not represent as such a regulatory mechanism. Instead, they could reflect long-range secondary structure constraints, which could be more efficiently captured by a different type of representation than the CNN allows.

A main application of DL models trained on synthetic sequences is the design of constructs with desired translation outputs [14]. While this has been demonstrated within the setting of the MPRA, where randomized 5’UTRs drive the expression of proteins such as eGFP or mCherry, whether the same accuracy can be achieved for endogenous mRNAs of interest remains to be determined. [14] have tested the same 5’UTR library in the context of the eGFP and mCherry coding regions and found that the model trained on the eGFP constructs explains 77-78% of the variance in mCherry expression, contrasting the 93% of the variance explained in eGFP expression. Interestingly, this difference has been attributed, in part, to differences in the polysome profiling protocol [14]. This points to the importance of establishing robust experimental protocols for generating reference data sets. However, the different coding regions likely contribute to the discrepancy in prediction accuracy as well, underscoring the importance of measuring the same library of constructs in different systems to identify the mechanisms responsible for a specific readout.

## 4 Conclusion

In summary, our analysis suggests a few directions for the study of translation control and applications to protein expression. First, to continue to uncover mechanisms that explain the expression of endogenous sequences, it will be important to include these sequences in high-throughput assays. The method of choice for measuring the translation output of endogenous mRNAs is ribosome profiling, a method that, on its own, is very reproducible (*R*^2^ ≳ 0.8). However, by factoring in the mRNA-seq-based estimation of mRNA abundance to calculate the TE adds leads to increased error in the estimate. Ensuring high accuracy of mRNA-seq and ribo-seq is important for obtaining reference data sets of TE. An additional limitation of endogenous mRNA translation data is its size. Currently, the number of mRNAs whose TE is estimated in a typical experiment is *∼* 20*^′^*000, which corresponds roughly to one isoform per gene. Accurate estimation of the TE of individual isoforms could be an important direction of methodological development [35, 36]. However, it is unlikely that many isoforms are simultaneously expressed at a high enough level to be accurately measured in a given cell type, or that sufficiently accurate data can be currently obtained from single cells [37] that express distinct isoforms. As a suboptimal alternative, TE measurements could be obtained in closely related cell types in which sufficient variation of transcription and thereby translation start sites occurs. In terms of training DL models on such data, an important consideration will be to ensure that training and test sets do not contain related sequences, to prevent models from achieving high prediction accuracy simply based on sequence similarity, without learning the regulatory grammar [38].

Second, towards predicting the impact of SNPs on translation, accurate models of endogenous mRNA expression are needed. As we have seen here, architectures beyond CNNs are desirable, and models used in natural language processing may provide a useful stepping stone. However, it will be interesting to develop architectures that can represent long-range dependencies of RNA secondary structures, perhaps also incorporated co-evolution constraints, as done for protein structure prediction [39, 40].

Third, towards the goal of designing constructs with specified translation outputs, it will be important to first determine the range of variation afforded by randomized 5’UTR variants by actually measuring the range of protein expression that can be covered with these variants. If this is sufficient, it will be important to determine the impact of unexplored parameters, such as the cellular context of construct expression and the impact of the coding region downstream of the randomized construct. For the former, the same construct library can be tested in various cell types, especially those that are closest to the cell type in which the mRNAs will be ultimately expressed (e.g. muscle cells for mRNA vaccines) [41]. Regarding the coding region, it will be interesting to test at least a few that cover the range of endogenous expression, from mRNAs with different life times and codon bias.

To conclude, DL models can be trained to very high precision on synthetic data, irrespective of their architecture. However, so far, synthetic data does not appropriately cover the space of regulatory elements influencing translation initiation. To achieve a comprehensive and predictive model as well as understand translation, training on endogenous sequences is necessary. The main bottleneck at the moment is obtaining sufficient and highly reproducible data on the translation of endogenous mRNAs. Experiments in a single cell type such as a cell line may not yield sufficiently many reliably measured 5’UTRs to train models such as TranslateLLM can be retrained. Perhaps this limitation can be circumvented by collecting data from multiple cell types, as they may contain distinct isoforms, with distinct 5’UTRs and translation efficiencies. Such a model could then be used for a broad variety of tasks, such as predicting the effect of point mutations, the translation efficiency of synthetic constructs, and for deepening our mechanistic understanding of translational control.

## 5 Methods

### 5.1 Experimental methods

We outline the experimental procedure for RNA and ribosome footprint sequencing of HepG2 cells.

#### 5.1.1 Cell culture

The HepG2 cell line was obtained from the laboratory of Dr. Salvatore Piscuoglio (DBM, Basel) and was cultured in Dulbecco’s Modified Eagle Medium (DMEM) containing 4.5 g/l glucose, 10% fetal calf serum, 4 mM L-glutamine, 1X NEAA, 5 ml Penicillin-Streptomycin at 5% *CO*_2_, 37*^◦^*C. Cells were passaged every 3-4 days.

#### 5.1.2 Cell lysis and collection

Cells were grown in 15 cm dishes to achieve a 70-80% confluency. Medium was replenished three hours prior to cell lysis. Cycloheximide (CHX) was added to a final concentration of 100 *µ*g/ml to arrest elongating ribosomes. Medium was immediately discarded and cells were washed once with ice-cold PBS containing 100 *µ*g/ml CHX. 500 *µ*l lysis buffer (20 mM Tris-HCl pH 7.5, 100 mM NaCl, 10 mM MgCl_2_, 1% Triton X-100, 2 mM dithiothreitol (DTT), 100 *µ*g/ml CHX, 40 U/*µ*l RNasin plus RNase inhibitor (Promega), 2 U/*µ*l Turbo DNase (Invitrogen) and EDTA-free protease inhibitor cocktail (Roche)) was added directly to the cells on the Petri dish. Cells were scraped and collected into 1.5 ml Eppendorf tubes. Then, samples were incubated for 5 min at 4*^◦^*C at continuous rotation (60 rpm), passed through a 23G needle for 10 times, and again incubated for 5 min at 4*^◦^*C at continuous rotation (60 rpm). Lysates were clarified by centrifugation at 3’000 xg for 3 min at 4*^◦^*C. Supernatants were centrifuged again at 10’000 xg for 5 min at 4*^◦^*C.

#### 5.1.3 Ribosome footprint sequencing

The ribosome footprinting sequencing protocol was adapted from protocols described in Refs. [18, 42, 43]. An equivalent to 8 OD260 of lysate was treated with 66 U RNase I (Invitrogen) for 45 min at 22*^◦^*C in a thermomixer with mixing at 1000 rpm. Then, 200 U SuperaseIN RNase inhibitor (20 U/*µ*l, Invitrogen) was added to each sample. Digested lysates were loaded onto 10-50% home-made sucrose density gradients in open-top polyclear centrifuge tubes (Seton Scientific). Tubes were centrifuged at 35’000 rpm for 3 hours at 4*^◦^*C (SW-41 Ti rotor, Beckmann Coulter ultracentrifuge. Samples were fractionated using the Piston Gradient Fractionator (Biocomp Instruments) at 0.75 ml/min by monitoring A260 values. Thirty fractions of 0.37 ml were collected in 1.5 ml Eppendorf tubes, flash frozen, and stored at *−*80*^◦^*C. The fractions (typically 3 or 4) corresponding to the digested monosome peak were pooled. RNA was extracted using the hot acid phenol/chloroform method. The ribosome-protected RNA fragments (28-32 nt) were selected by electrophoresis on 15% polyacrylamide urea TBE gels and visualized with SYBR Gold Nucleic Acid Gel Stain (ThermoFisher Scientific). Size selected RNA was dephosphorylated by T4 PNK (NEB) for 1 h at 37*^◦^*C. RNA was purified using the acid phenol/chloroform method. Depletion of rRNA was performed using the riboPOOL kit (siTOOLs biotech) from 433 ng of RNA according to the manufacturer’s instructions. Libraries were prepared using the SMARTer smRNA-Seq Kit for Illumina (Takara) following the manufacturer’s instructions from 15 ng of RNA. Libraries were purified by electrophoresis on 8% polyacrylamide gels and sequenced on the Illumina NextSeq 500 sequencer in the Genomics Facility Basel (Department of Biosystems Science and Engineering (D-BSSE), ETH Zürich).

#### 5.1.4 RNA-sequencing

RNA was extracted from 15 *µ*l of cell lysate using the Direct-zol RNA Microprep Kit (Zymo Research) following the manufacturer’s instructions and including DNAse treatment for 15 min at room temperature. Samples were eluted with 15 *µ*l nucleasefree water. The RNA integrity numbers (RIN) of the samples were between 9.9 and 10.0, measured using High Sensitivity RNA ScreenTape (TapeStation system, Agilent). RNA was quantified using a Qubit fluorometer. Libraries were prepared using the SMART-seq Stranded for total RNA-seq kit (Takara) from 5 ng of RNA and sequenced on the Illumina NextSeq 500 sequencer in the Genomics Facility Basel (Department of Biosystems Science and Engineering (D-BSSE), ETH Zürich).

### 5.2 Data sets

The data sets used in this study are as follows.

#### 5.2.1 Optimus50

Constructs consisted in 25 nts of identical sequence (for PCR amplification) followed by a 50 nt-long random 5’UTR sequence upstream of the GFP coding region. Their sequences and associated mean ribosome load measurements were obtained from the GEO repository, accession number GSE114002. Non-sequential features were computed and annotated for each sequence with a python script. The normalized 5’UTR folding energy was determined with the RNAfold program from the ViennaRNA package [44]. The G/C-fraction was calculated using the biopython package [45]. Number of OOF and IF uAUGs were calculated with standard python methods. ORF/UTR length and number of exons were identical in this data set and therefore uninformative. Following [14], we split the 20’000 5’UTRs with the highest coverage in mRNA seq for testing and kept the rest for training.

#### 5.2.2 Optimus100

Constructs were made from random sequences, human 5’UTRs of suitable size (25-100 nts), their single nucleotide polymorphism-containing variants and 3’-terminal fragments of longer 5’UTRs. MRL measurements were done as for the Optimus50 data set. Sequences and associated MRL estimates were obtained from the GEO repository, accession number GSE114002. The non-sequential features were computed just as for Optimus50, with the UTR length being an additional degree of freedom. The 5’000 5’UTRs with highest coverage in mRNA-seq are held out for testing, just as in [14].

#### 5.2.3 Human genetic variants

Sample *et. al.* [14] extracted 3’577 5’UTR SNPs from the ClinVar database [29] and constructed variant 5’UTRs containing these SNPs. These variants were transfected to HEK293 cells and the respective MRL was measured as described in the paragraph about Optimus50. We also appended non-sequential features as outlined there, with the UTR length is an additional variable. The sequences and MRL were downloaded from GEO repository GSE114002.

#### 5.2.4 Yeast50

Yeast colonies were grown in media without *HIS3*. Yeast cells were transduced with plasmids containing the *HIS3* -ORF attached to a random pool of *∼* 500*^′^*000 randomized 50*nt* long 5’UTRs. The growth rate is directly controlled by the amount of *HIS3* protein, which only is controlled by the 5’UTR sequence. The data were obtained from GEO, accession number GSE104252. The calculation of non-sequential features followed the exact same procedure as for Optimus50. The top 5% 5’UTRs in terms of read coverage were used for testing.

#### 5.2.5 DART

We downloaded the training data from Supp. Tab. S2 of [15]. Non-sequential features were calculated as for Optimus50. Since uAUGs are mutated in this data set to avoid ambiguity in the translation start site, we did not include the number of OOF or IF uAUGs in the list of non-sequential features to learn from. Also, DART uses a luciferase reporter only including the first bit of the coding sequence, so neither the number of exons nor the CDS length are meaningful, therefore, we did not include these features, either. The first bit of the CDS sequence is available as a separate column in their Supp. Tab. S2. We use three different random splits of 10% of the data for testing.

#### 5.2.6 Human mRNA sequences

The human transcript sequences were pulled from ENSEMBL [46] with pybiomart. We use the latest annotation (GRCh38.105) and the latest sequence file. The human transcriptome sequences were assembled with gffread [47].

#### 5.2.7 Yeast mRNA sequences

We used the current release of the yeast genome R64.1.1 [48] with the current annotation R64.1.1.110 from the Saccharomyces cerevisiae Genome Database (SGD). We enriched this annotation with the longest annotated transcript from TIF-seq, see [49], providing us with 5’UTR sequences. Gffread [47] yielded the yeast transcriptome.

#### 5.2.8 Yeast TE data

Following [6], we downloaded the RNA-seq data from the SRA archive (SRR2986251), and ribosome-profiling data from an older archive (SRR1049521). The riboseq analysis was conducted as in [50], the RNA-seq was performed using zarp [51]. All nonsequential features (log ORF length, UTR length, G/C-content fraction of the UTR, number of exons, number of OOF uAUGs, number of IF uAUGs, normalized 5’UTR folding energy) were computed or extracted from the genome annotation. The 10% of transcripts with the highest TIN were used for testing purposes.

#### 5.2.9 HEK 293 TE data

Ribo-seq and mRNA-seq data were obtained from the European Nucleotide Archive, accession PRJNA591214. The riboseq analysis was conducted as in [50], the RNA-seq was performed as in [51]. For the calculation of the translation efficiency, we only took into account RNA-seq and ribo-seq reads in the CDS, not on the entire transcript. For stringency in the attribution of reads to mRNAs, we calculated relative isoform abundances by running salmon [52] on the RNA-seq samples and selected the most abundant isoform as representative, to which we mapped the RNA and ribo-seq reads. The 10% of transcripts with the highest TIN (squared average over the three replicates) were used for testing.

#### 5.2.10 HepG2 TE data

We followed the experimental procedure outlined in the experimental methods. The rest of the analysis was done as for the HEK293 TE data.

#### 5.2.11 ClinVar data

We downloaded the most recent version of the ClinVar data base vcf file (vcf GRCh38 [30]). With bedtools-intersect [53], we identified variants from ClinVar in annotated genes and only kept variants of annotated 5’UTRs. With a python script, we calculated the coordinates of the polymorphisms on all affected transcripts. Then, we constructed the variant 5’UTRs in the human transcriptome (created with gffread [47] from the current ENSEMBL annotation GRCh38.105) and extracted the coding regions. This left us with 84’127 mutated transcripts. Next, we computed the non-sequential features as for Optimus50. We predicted the variant TE with the transfer-learning version of TranslateLLM (trained on human endogenous HEK 293, pre-trained on the Optimus100 data set). Matching the transcript variants and predictions to transcripts for which we have TE measurements left us with 7’238 transcripts.

### 5.3 Model architectures

We implemented previous published models that predict the translation output from 5’UTR sequences according to the their description in the respective studies.

#### 5.3.1 Optimus5’

Optimus5’ was the first neural network trained to predict translation initiation efficiency [14]. It consist of three convolutional layers with 120 8nt-long filters each. They all feature a relu activation function and are succeeded by two dense layers, one reducing the input dimensionality to 40 with another relu nonlinearity, and a last dense layer reducing to a single number. The two last layers are separated by a dropout layer that stochastically ignores 20% of the input signals during training. The configuration allowing predictions for 5’UTR s up to 100 nts in length has 714’681 parameters. Two different configurations of Optimus5’ were proposed: one trained of a pool of *∼* 280*^′^*000 5’UTR sequences of 50 nts, and another trained on a pool of *∼* 105*^′^*000 5’UTRs of 25 *−* 100 nts. Variable lengths were handled by anchoring the 5’UTR at their 3’end (adjacent to the start codon) and padding the 5’ end with 0’s in the one-hot encoded representation. Since endogenous 5’UTR s vary widely in length, we used the latter configuration and data set, considering it to be more realistic. However, the size of the model is also larger. To run the Optimus models on the local scientific computing infrastructure, the model training was re-implemented in a python script, rather than a jupyter notebook as in the git repository cited in [14].

#### 5.3.2 FramePool

FramePool [16] technically overcomes the limitation on 5’UTR length. While also relying on convolutional layers and a final two-layer perceptron, FramePool introduced an operation called ’framewise pooling’. This was motivated by previous observations that out-of-frame uORFs have a strong impact on the translation output. Framewise pooling involves the pooling of output of convolutional layers separately for the +0, +1, and +2 frames. The subsequent multi-layer perceptron (MLP) takes as input the average and the maximum of each of the three pools (per convolutional filter). This makes the input of the final MLP independent of the input length and allows for varying UTR lengths from a technical standpoint. Trained on the same data sets as Optimus5’, the performance on data of varying UTR length was increased. The number of parameters in Framepool is only about a third of what Optimus5’ requires for UTR lengths *≤* 100nt, namely 282’625 parameters. We pulled Framepool from the git repository referenced in [16]. A python script related the model to our format of input data.

#### 5.3.3 MTtrans

While CNNs are generally not a natural choice when it comes to modeling sequences of variable length, recurrent neural networks (RNNs) were developed exactly for this purpose. Conventional RNNs suffer from the so-called vanishing gradient problem, whereby memory of distant context is lost. Moreover, they can only memorize the leftside context, since they process sequences from left to right. These problems are solved by long-short term memory units (LSTM) [24] and bidirectional layers. However, as there is no correspondence between output cells and position in the sequence, the interpretability of this type of model is more challenging. MTtrans [17] has been proposed as an attempt to get the best of both CNN and LSTM worlds. It follows the general idea of detecting motifs, with four convolutional layers stacked on top of each other, batch normalization, L2 regularization and dropout layers in between to avoid over-fitting and ensure stability. This component of MTtrans is called ’shared encoder’ and is followed by two bidirectional gated recurrent unit (GRU) [54] layers and two dense layers to make the final prediction. GRUs are quite similar to LSTMs, but they do not feature an output-gate [24, 54], and therefore have fewer weights to adjust than LSTM layers. This second component of MTtrans is called ’task-specific tower’, because it is re-trained for each data set (task), while the encoder is shared across all tasks. By training the encoder on data sets of different organisms and cells, the authors aim to capture general features of translation that apply to all of the studied systems. This is an example of transfer learning, hence the ’trans’ in the name MTtrans. MTtrans appears to be considerably bigger than its two predecessors, with *∼* 2*^′^*100*^′^*000 parameters. A re-evaluation of the results in [17] was unfortunately not possible since the code in the provided github repository was still work in progress. Therefore, we attempted reconstructing MTtrans in our own ML framework, but will quote the numbers reported in [17], wherever available.

#### 5.3.4 TranslateLLM

The start of the coding region has a biased nucleotide composition that also plays a role in translation initiation (*c.f.* [55]). Putting the first 100nts into another bidirectional LSTM model therefore provides additional information about initiation likelihood. These three models, bidirectional LSTM for 5’UTRs, bidirectional LSTM for beginning of ORF, and non-sequential features, can now be concatenated into a big model.

There is, of course, a lot of redundance in these inputs, as the folding energy of the 5’UTR is determined by its nucleotide sequence, GC-content and length of the 5’UTR. One way to mitigate this redundance is to use high dropout rates after the final bidirectional LSTM layer of both RNNs (5’UTR and ORF). For training from scratch, we used dropout rates of 0.8 for the 5’UTR model and 0.2 for the CDS model. After concatenating the numerical input data with the two bidirectional LSTMs, a final dense layer computes a single number, the logarithm of the translation efficiency, scaled to a Gaussian distribution with unit variance and expectation value 0. The network was implemented in python, using the keras API with a tensorflow backend [56, 57]. We used the adam algorithm for training [58]. Beyond dropout layers, overfitting is prevented by imposing an early stopping criterion. Of the randomly reshuffled training data, 20% serve validation purposes. To further improve the performance of the model, we pretrained it on the variable-length Optimus100 data set before training on the endogenous data. In that scenario, we used slightly lower dropout rates for thr 5’UTR LSTM, of 0.5.

### 5.4 Linear models

As the translation initiation efficiency was reported to be explained, to a large extent, by a small number of mRNA features [6], we have included in our study two variants of a small linear model. The features were as follows. First, upstream open reading frames (uORFs) were reported in many studies to reduce the initiation of translation at the main ORF [59]. The effect was found to be largely due to uORFs that are in a different frame than the main ORF, which we have referred to as ”out-of-frame ORFs” or ”out-of-frame AUGs”, because the presence and position of stop codons matching these ORFs is not generally considered. Thus, one of the linear models included the numbers of out-of-frame and in-frame AUGs, while the other only the former. The secondary structure of 5’ cap-proximal region of the mRNA is known to interfere with the binding of the eIF4F cap-binding-complex [5], and thus a weak positive correlation has been observed between the free energy of folding of the first 80 5’UTR nts and the translation initiation efficiency of yeast mRNAs [6–8]. A more minor impact on yeast translation has also been attributed to the ORF length (negative correlation with TIE) [6, 60], 5’UTR length and G/C content [6]. For human cells, the number of exon-exon junctions has also been linked to TIE [22]. Suppl. Fig. 3 shows density plots of these parameters, comparing the major data sets we used, *i.e.* the three MPRA data sets, DART, and the two endogenous ones.

The linear models are of compelling simplicity: They only only have as many parameters as features they cover, plus a single global bias term. For instance, the linear model describing the Optimus50 data set consists of weights multiplying the normalized 5’UTR folding energy, the G/C-content, the number of IF and OOF upstream AUGs, and the bias term, totaling to 5 parameters.

## Declarations

### Ethics approval

Not applicable.

### Consent to participate

Not applicable.

### Consent for publication

Not applicable.

### Availability of data and materials

Sequencing data from ribosome footprinting and RNA sequencing in the HepG2 cell line are available under BioProject ID PRJNA1045106. The evaluated clinical variants from the clinvar data base are attached in Supp. Tab. 1. MPRA measurements in HEK293 cells from [14] are publicly available from GEO repository GSE114002, MPRA measurements in yeast (see [20]) GSE104252. DART measurements from yeast data are available from Supp. Tab. 2 of [15]. Yeast RNA sequencing and ribosome footprinting data from [6] were retrieved from the SRA under accession numbers SRR2986251 and SRR1049521. RNA sequencing and ribosome footprinting data from HEK293 cells (c.f. [19]) for this study were downloaded from the European Nucleotide Archive under accession number PRJNA591214.

### Code availability

The code for TranslateLLM and data-preprocessing is publicly available under https://git.scicore.unibas.ch/zavolan_group/data_analysis/predicting-translation-initiation-efficiency.

### Authors’ contributions

M.Z. and N.S. conceived the study. N.S. implemented the models and carried out the analysis. M.P. and A.G. provided experimental data.

## Acknowledgments

We would like to thank Aleksei Mironov for providing us with yeast 5’ UTR annotations and, along with Meric Ataman, for helpful discussions.

