## Supplementary Information for "Current limitations in predicting mRNA translation with deep learning models"

### 1 Supplementary Figures

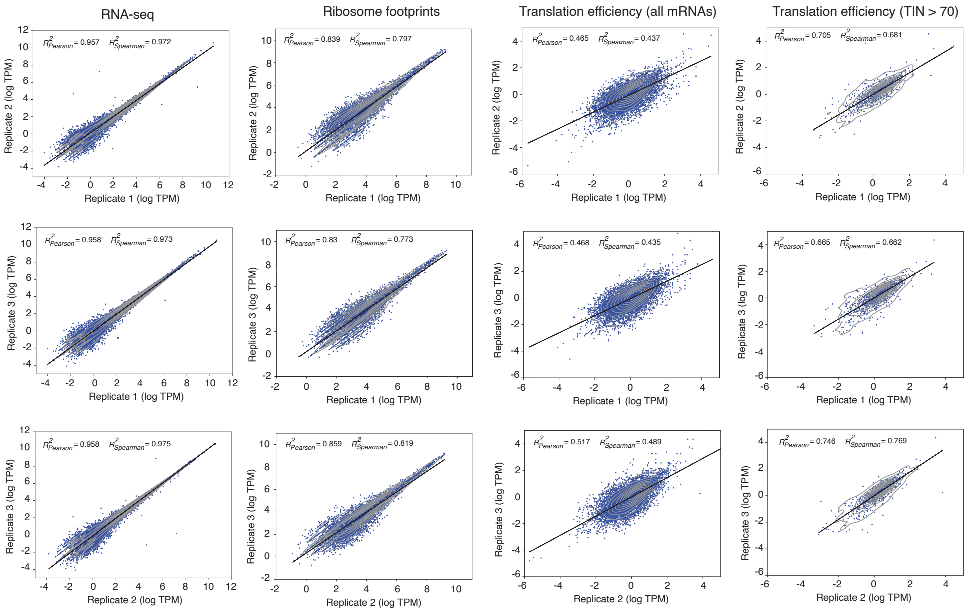


**Fig. 1** Inter-replicate reproducibility of different experiments in HEK 293 cells: mRNA-sequencing coverage of the CDS (col. 1), ribosome footprint coverage of the CDS (col. 2), translation efficiency (TE) as measured by the ratio of the former two (col. 3), and TE of transcripts with TIN ¿ 70 (col. 4).

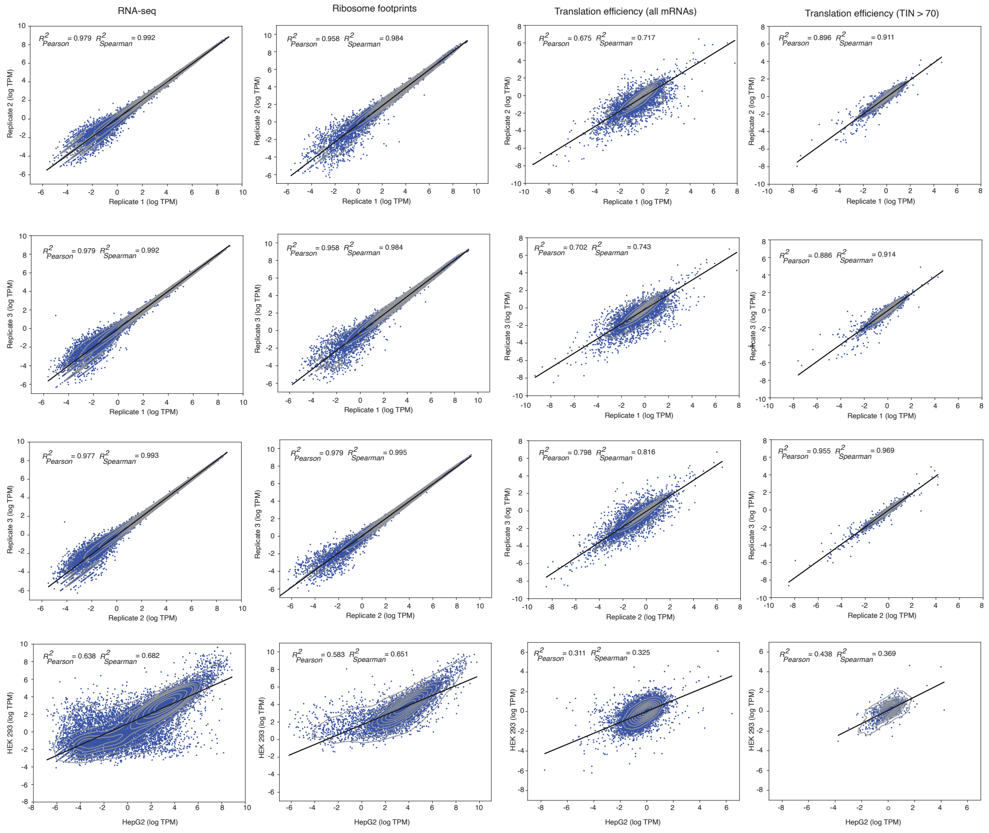


**Fig. 2** Inter-replicate of HepG2 cell line and inter-tissue correlation between HEK 293 and HepG2 cell line: mRNA-sequencing coverage of the CDS (col. 1), ribosome footprint coverage of the CDS (col. 2), translation efficiency (TE) as measured by the ratio of the former two (col. 3), and TE of transcripts with TIN $> 70$ (col. 4). Correlation between replicates 1 and 2 in line 1, replicates 1 and 3 in line 2, 2 and 3 in line 3, and HepG2 and HEK293 in line 4.

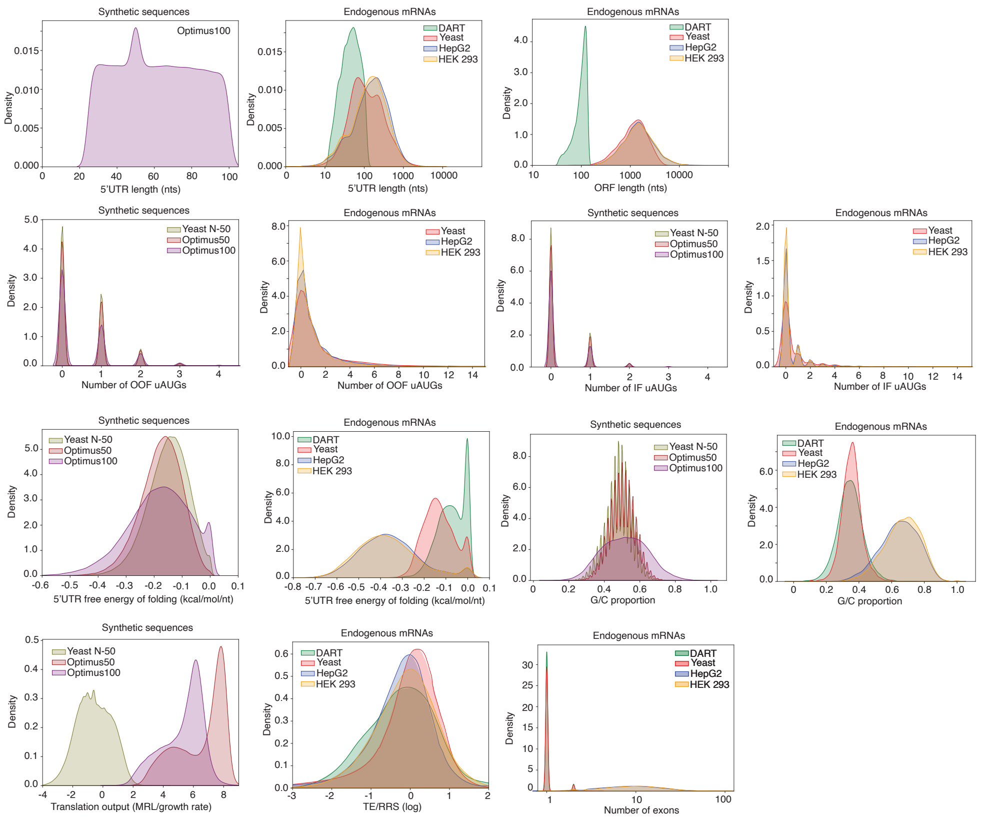


**Fig. 3** Density plots of non-sequential features in the six different data sets used in this study. Synthetic data sets are Yeast50 [1], Optimus50, and Optimus100 [2]. Endogenous data sets are DART [3], endogenous yeast [4], and endogenous HEK293 [5]. Non-sequential features are the 5' UTR length (cols. 1,2, l. 1), the mORF length (col. 3, l. 1), the number of out-of-frame (OOF) uAUGs (cols. 1,2, l. 2), the number of in-frame (IF) uAUGs (cols. 3,4, l.2), the 5'UTR folding energy per unit base (col2. 1,2, l. 3), the G/C-content fraction (cols. 3,4, l. 3), and the number of exons (col. 3, l. 4). The translation output as measured by mean ribosome load (MRL) or yeast growth rate (col. 1, l. 4), and log ribosome recruitment score (RRS) or log translation efficiency (TE) (col. 2,l. 4) is also shown.

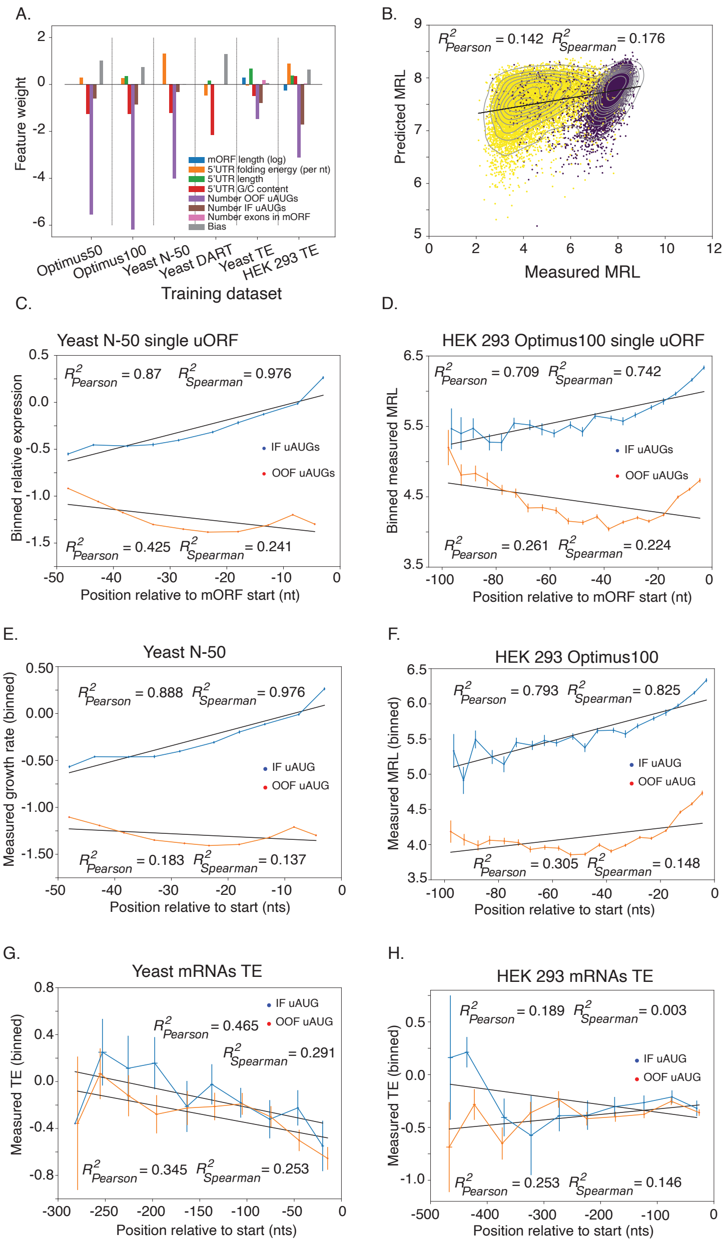


**Fig. 4** Weights of the linear model associated with the different non-sequential features when being trained on different data sets (A). Upstream AUGs are by far the most influential feature. When training Optimus5' on data lacking uAUGs, it loses most of its predictive power (B). The distance of IF and OOF uAUGs to the start of the mORF determines the inhibitory strength, as can be seen for yeast MPRA data (C, E), and HEK 293 MPRA data (D, F). Endogenous data yeast data are shown in (G), HEK 293 data in (H). Panels (C) and (D) display only data from transcripts with a single uAUG, whereas (E-H) use data from transcripts with at least one uAUG (either only IF or OOF). The computation of the position is based on the most upstream AUG.

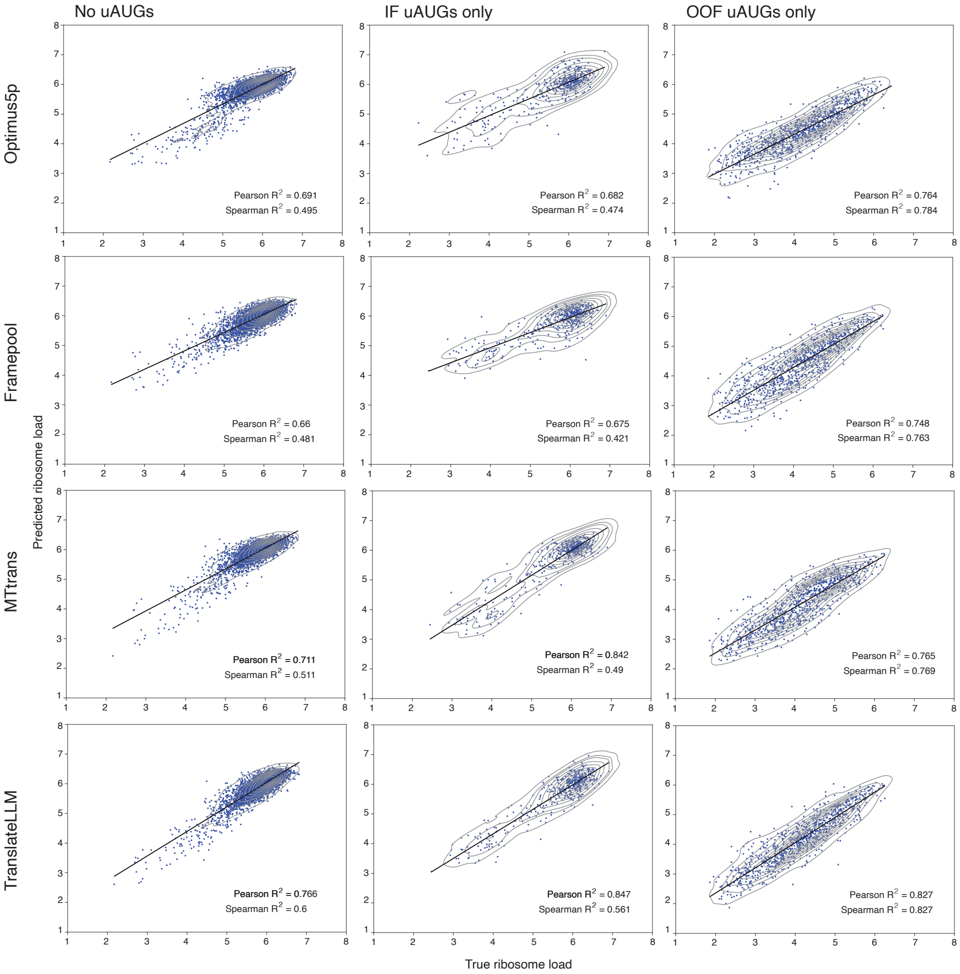


**Fig. 5** The Optimus100 data set was split into three subcategories: 5'UTRs containing no uAUGs at all, 5'UTRs containing at least one IF uAUG, but no OOF uAUGs, and 5'UTRs containing at least one OOF uAUG, but no IF uAUGs. We evaluate the performance of the three existing deep learning architectures (Optimus5', FramePool, MTtrans), as well as the new architecture, TranslateLLM, on the three data sets. TranslateLLM predicts all three subsets with the highest accuracy and leads to the most evenly shaped contour plots.

#### 2 Supplementary Tables

Supplementary Table containing clinvar 5'UTR variants [6], their predicted log-fold change in TE (LFC TE), clinical significance, mutated and parent UTR, gene and transcript ID. Attached to submission as `supp_tab_1.tsv`
